# Remote control of CAR T cell therapies by thermal targeting

**DOI:** 10.1101/2020.04.26.062703

**Authors:** Ian C. Miller, Lee-Kai Sun, Adrian M. Harris, Lena Gamboa, Ali Zamat, Gabriel A. Kwong

## Abstract

The limited ability to control anti-tumor activity within tumor sites contributes to poor CAR T cell responses against solid malignancies. Systemic delivery of biologic drugs such as cytokines can augment CAR T cell activity despite off-target toxicity in healthy tissues that narrows their therapeutic window. Here we develop a platform for remote control of CAR T therapies by thermal targeting. To enable CAR T cells to respond to heat, we construct synthetic thermal gene switches that trigger expression of transgenes in response to mild elevations in local temperature (40–42 °C) but not to orthogonal cellular stresses such as hypoxia. We show that short pulses of heat (15–30 min) lead to more than 60-fold increases in gene expression without affecting key T cell functions including proliferation, migration, and cytotoxicity. We demonstrate thermal control of broad classes of immunostimulatory agents including CARs, Bispecific T cell Engagers (BiTEs), and cytokine superagonists to enhance proliferation and cell targeting. In mouse models of adoptive transfer, photothermal targeting of intratumoral CAR T cells to control the production of an IL-15 superagonist significantly enhances anti-tumor activity and overall survival. We envision that thermal targeting could improve the safety and efficacy of next-generation therapies by allowing remote control of CAR T cell activity.

## INTRODUCTION

Engineered T cell therapies such as Chimeric Antigen Receptor (CAR) T cells are transforming clinical care for hematological malignancies, spurring numerous efforts to expand their use for different cancer types and applications. However, this success has not reliably translated to solid tumors^1^. The factors that contribute to low response rates are multifaceted and include the paucity of tumor-specific antigens, inefficient persistence and expansion of adoptively transferred T cells, and immunosuppression by the tumor microenvironment (TME)^2^. Promising approaches to improve anti-tumor activity of engineered T cells include systemic administration of potent immunostimulatory agents such as cytokines, checkpoint blockade inhibitor antibodies, and bispecific T cell engagers (BiTEs)^3-6^. However, these biologics lack specificity, activate both engineered and endogenous immune cells, and exhibit toxicity in healthy tissue which limits maximum tolerable doses and narrows their therapeutic windows^7-10^. Thus, expanding current abilities to target and locally augment CAR T cell functions at tumor and disease sites such as draining lymph nodes could improve the safety and efficacy of cell-based therapies.

Emerging strategies to control engineered T cells and augment their anti-tumor activity include the use of biomaterials to co-deliver adjuvants to the TME as well as genetic constructs for autonomous expression of immunostimulatory genes. For example, implantation of biopolymer scaffolds loaded with tumor-specific T cells and immunostimulatory adjuvants at the surgical site improved postoperative responses following primary tumor resection in mouse models^11, 12^. To provide a localized source of adjuvants, T cells tethered on their cell surface to nanoparticle ‘backpacks’ allowed infiltrating T cells to carry cargo^13^ and release a one-time dose of drug within tumors^14^. Increasingly sophisticated genetic circuitry has also allowed T cells to locally produce biologics to overcome immunosuppression or target antigens after tumor infiltration. For example, ‘armored CARs’ leverage constitutive expression of biologics such as IL-12^15^, αPD-1 scFvs^16^, and BiTEs^17^ to improve anti-tumor activity. T cells have also been engineered with sense-and-respond biocircuits that conditionally activate in the presence of specific input signals. These strategies include NFAT-inducible cassettes that upregulate expression of cytokines following T cell recognition of a tumor-associated antigen^18-20^. To further increase specificity, T cells have been engineered to target unique combinations of epitopes expressed in the TME to allow discrimination from healthy cells expressing a single epitope^21^. Such approaches based on Boolean logic require the presence of both target antigens for T cell activation to occur and have demonstrated efficacy in multiple models of focal tumors^22, 23^. Collectively, these approaches illustrate the need to develop strategies to control and improve intratumoral T cell activity.

Here we developed a platform for remote thermal control of CAR T cell therapies. Heat treatments are used clinically to sensitize cancer cells to chemotherapy, ablate isolated metastatic nodules, and enhance diffusion of small molecule drugs into tumors^24^. Both superficial and deep-seeded tumors can be targeted for thermal treatment by platforms including high intensity focused ultrasound (HIFU)^25^, laser interstitial thermal therapy (LITT)^26^, and electromagnetic heating^27^. To engineer T cells with the ability to respond to heat, we constructed and screened panels of synthetic thermal gene switches containing combinations of Heat Shock Elements (HSEs) and core promoters to identify an architecture that responds to mild hyperthermia while remaining non-responsive to orthogonal cell stresses. We designed thermal constructs to control broad classes of immunostimulatory genes including CARs, BiTEs, and a cytokine superagonist to enhance key T cell functions including proliferation and T cell targeting. In an adoptive transfer model of cancer, remote thermal control of an IL-15 superagonist improved anti-tumor CAR T cell activity by reducing tumor burden and improving survival of tumor-bearing animals. Remote control of CAR T cells by thermal targeting could improve the precision of cellular therapies by enabling site-specific control of anti-tumor responses.

## RESULTS

### Engineering thermal-specific gene switches

The cellular response to hyperthermia is mediated by trimerization of the temperature-sensitive transcription factor Heat Shock Factor 1 (HSF1) and its subsequent binding to DNA motifs termed Heat Shock Elements (HSEs). HSEs are comprised of multiple inverted repeats of the consensus sequence 5’-nGAAn-3’^28, 29^ and are arrayed upstream of HSPs thereby enabling their upregulation following thermal stress^30^. The response of endogenous HSP genes is selective, but not specific, for heat as their promoters contain additional regulatory elements (e.g., hypoxia response elements^31^, metal-responsive elements^32^) that mediate transcription following exposure to a diverse set of cues including hypoxia^33^, heavy metals^34^, and mechanical force^35^. Moreover, differences in the core promoter (e.g., initiator elements, TATA box) influence the composition of the pre-initiation complex (PIC) and its interactions with transcriptional enhancers including HSF1, providing an additional mechanism whereby transcriptional responses to heat are regulated differently across tissues and types of cells^36, 37^. Due to this complexity and cross-activation of HSP promoters by non-thermal response pathways, we sought to build synthetic gene switches that are activated by heat but not by other sources of stress.

To design thermal-specific gene switches, we cloned 6 candidate constructs comprised of 2 to 7 repeats of the HSE motif 5’-nGAAnnTTCnnGAAn-3’ upstream of the Hspb1 core promoter into Jurkat T cells (labeled 2H-B1 to 7H-B1, **Fig. 1a**). We initially selected the Hspb1 core promoter as its parent gene was one of two that were upregulated by more than 20-fold at 42 °C in primary murine T cells in contrast to more than 80 HSP and HSP-related genes that did not respond to heat treatment (**Fig. S1**). We reasoned that by selecting a core promoter from an endogenous gene with a high thermal response, it would facilitate transcriptional activity when integrated with HSE repeats. To quantify responses of our thermal switches, transduced Jurkat T cells were transiently heated to 3–5 °C above body temperature (i.e., 40–42 °C), a mild temperature range in contrast to those used for ablative therapies (>50 °C)^25^. Compared to control samples at 37 °C, we observed increased expression of the reporter Gaussia luciferase (Gluc) as the temperature and number of HSEs increased (**Fig. 1b**). Constructs containing 5–7 HSE repeats (5H-B1 to 7H-B1) resulted in significantly higher thermal responses compared to those with 2–4 HSEs (2H-B1 to 4H-B1) (**Fig. 1c**). Collectively, these data confirmed that our synthetic thermal gene switches were heat-sensitive and that the magnitude of the response was dependent on HSE number.

**Figure 1.**
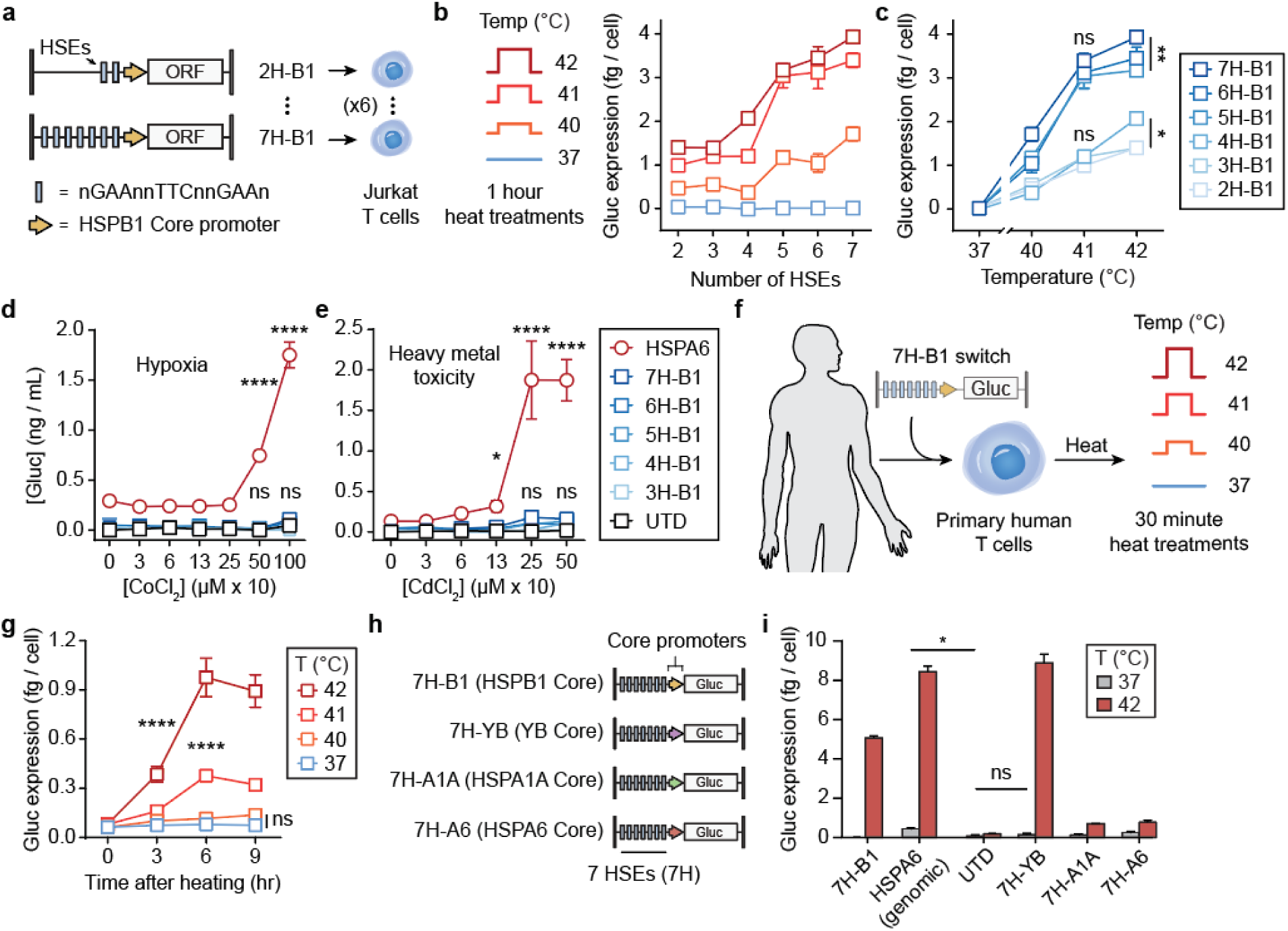
Constructing thermal-specific gene switches. (**a**) Schematic of B1 synthetic thermal gene switches. A panel of six constructs were cloned comprising 2 to 7 heat shock elements (HSEs) coupled to the HSPB1 core promoter into Jurkat T cells (labeled 2H-B1 to 7H-B1). Capitalized base pairs within HSE were conserved while base pairs indicated as n were randomized to facilitate synthesis. (**b**) Gluc reporter expression by Jurkat T cells following heating as a function of number of HSE or (**c**) temperature (ns = not significant, *P<0.05, **P<.01, two-way ANOVA and Tukey post-test and correction, error bars show SEM, n = 3). (**d**) Activity of synthetic thermal gene switches compared to endogenous HSPA6 promoter following exposure to CoCl_2_ to mimic hypoxia or (**e**) to CdCl_2_ to model heavy metal toxicity (ns = not significant, *P<0.05, ****P<.0001, two-way ANOVA and Tukey post-test and correction, error bars show SEM, n = 3). (**f**) Schematic for 30 minute thermal treatments of engineered primary human T cells at treatments ranging from 37 to 42 °C. (**g**) Kinetics of Gluc reporter expression by primary human T cells following heat treatments at indicated temperatures (****P<.0001, two-way ANOVA and Tukey post-test and correction, error bars show SEM, n = 3). (**h**) Schematic of core promoter screen using 7 HSE repeats. (**i**) Activity of thermal gene switches containing different core promoter constructs following heat treatments in primary human T cells (ns = not significant, *P<0.05, one-way ANOVA and Tukey post-test and correction, error bars show SEM, n = 3).

We next tested thermal specificity using hypoxia and heavy metal toxicity as two representative non-thermal stresses^38-40^. As a benchmark, we compared with the endogenous HSPA6 promoter, which is a highly stress-inducible HSP promoter^41^ previously used for thermal control of gene expression^42-45^. We tested the gene switches by incubating transduced Jurkat T cells with the hypoxia-mimetic agent CoCl_2_ – a stabilizer of the hypoxia response’s master regulator Hypoxia Inducible Factor-1α (Hif-1α)^46^ – as well as the heavy metal complex cadmium chloride (CdCl_2_). Whereas Jurkat T cells cloned with the HSPA6 promoter showed dose-dependent activation by hypoxia (**Fig. 1d**) and cadmium toxicity (**Fig. 1e**), T cells transduced with our synthetic gene switches (3H-B1 to 7H-B1) were not activated and remained statistically identical to untransduced (UTD) controls up to concentrations above the ranges commonly used to test cellular responses to hypoxia and cadmium^46-49^ (1000 µM CoCl_2_ and 500 µM CdCl_2_) (**Fig. 1d, e**). These results confirmed that our synthetic gene switches have increased thermal-specificity compared to endogenous HSPs and are not cross-activated when exposed to non-thermal stresses.

To test whether these constructs work in primary human T cells, we transduced T cells with the 7H-B1 thermal switch (**Fig. 1f**). Above a threshold temperature of 40 °C, we observed thermal switch activation that peaked between 3 to 6 hours after heating but was ∼40% less responsive compared to the endogenous HSPA6 (**Fig. 1g**,**h**). Because the Hspb1 core promoter was initially selected from a screen of murine T cells, we reasoned that thermal responses in primary human T cells may be further tuned with different core promoters. Thus, we compared additional core promoters including ones cloned from the HSPA1A gene identified in the qPCR screen (**Fig. S1**), the human HSPA6 gene, and a synthetic core promoter (YB) described previously^50, 51^ (**Fig. 1h**). Consistent with our previous observations, we observed negligible activation with heat treatments at 37–40 °C for 24 hours across all 4 constructs tested (**Fig. S2**). By contrast, after 30 minutes at 42 °C, the 7H-YB construct produced the highest Gluc reporter levels while maintaining statistically identical basal activity at 37 °C compared to untransduced controls. This corresponded to a ∼60-fold increase in thermal switch activity compared to ∼20-fold increases by the endogenous HSPA6 (**Fig. 1i**). On the basis of these data, we selected 7H-YB for subsequent experiments due to its improved thermal specificity and high thermal response.

### Primary T cells maintain key functions after thermal treatments

Next, we sought to identify the range of thermal delivery profiles that would be well-tolerated by primary T cells without affecting key functions including proliferation, migration, and cytotoxicity. In thermal medicine, heating target sites to temperatures greater than 50 °C is used to locally ablate tissue by inducing tumor cell apoptosis and coagulative necrosis^25^. By contrast, mild hyperthermia therapy (40–42 °C) is used to enhance transport of small molecules such as in Hyperthermic Intraperitoneal Chemotherapy (HIPEC) where abdominal infusions of heated chemotherapy serve as adjuvant treatment following surgical debulking in advanced ovarian cancer patients^24, 52, 53^. At temperatures below 45 °C, transient exposure to mild hyperthermia is well-tolerated by cells and tissues due to induction of stress-response pathways including HSPs^54^. In addition to temperature range, we also considered T cell responses to continuous and fractionated heat treatments. Dose fractionation is a commonly used approach in radiotherapy to reduce damage to normal tissues^55^ while maximizing the effect of radiation on cancer. Based on our previous observations that thermal pulse trains increased Jurkat T cell tolerance compared to continuous heat treatments with an identical treatment area-under-the-curve (AUC)^56^, we sought to additionally probe the effect of thermal dose fractionation on primary T cells.

We first compared pulsed heat treatments at 67% duty cycles comprised of three discrete thermal pulses (5 or 10 min each) separated by intervening rest periods at 37 °C (2.5 or 5 min each) to their unfractionated counterparts (15 or 30 min continuous heating) (**Fig. 2a**). Across the conditions tested, we observed significant increases in Gluc reporter expression with pulsed heat treatments at 42 (30 min AUC) and 43 °C (15 and 30 min AUC) by up to ∼30% compared to continuous delivery. To assess T cell viability, we quantified death (PI) and apoptotic (Annexin V) markers and observed no significant difference in viability between unheated, pulsed, and continuously heated samples for up to 30 minutes (**Fig. 2b**). By contrast a ∼34% reduction in T cell viability was observed in positive control samples that were continuously heat treated for 60 minutes. Similar trends were observed in T cell proliferation assays by dye (CTV) dilution where the percent of proliferated T cells following incubation with CD3/28 beads was unaffected by both continuous and pulsed heating for 30 minutes at 42 or 43 °C while positive control samples that were heated for 60 minutes resulted in reduced T cell proliferation (**Fig. 2c**).

**Figure 2.**
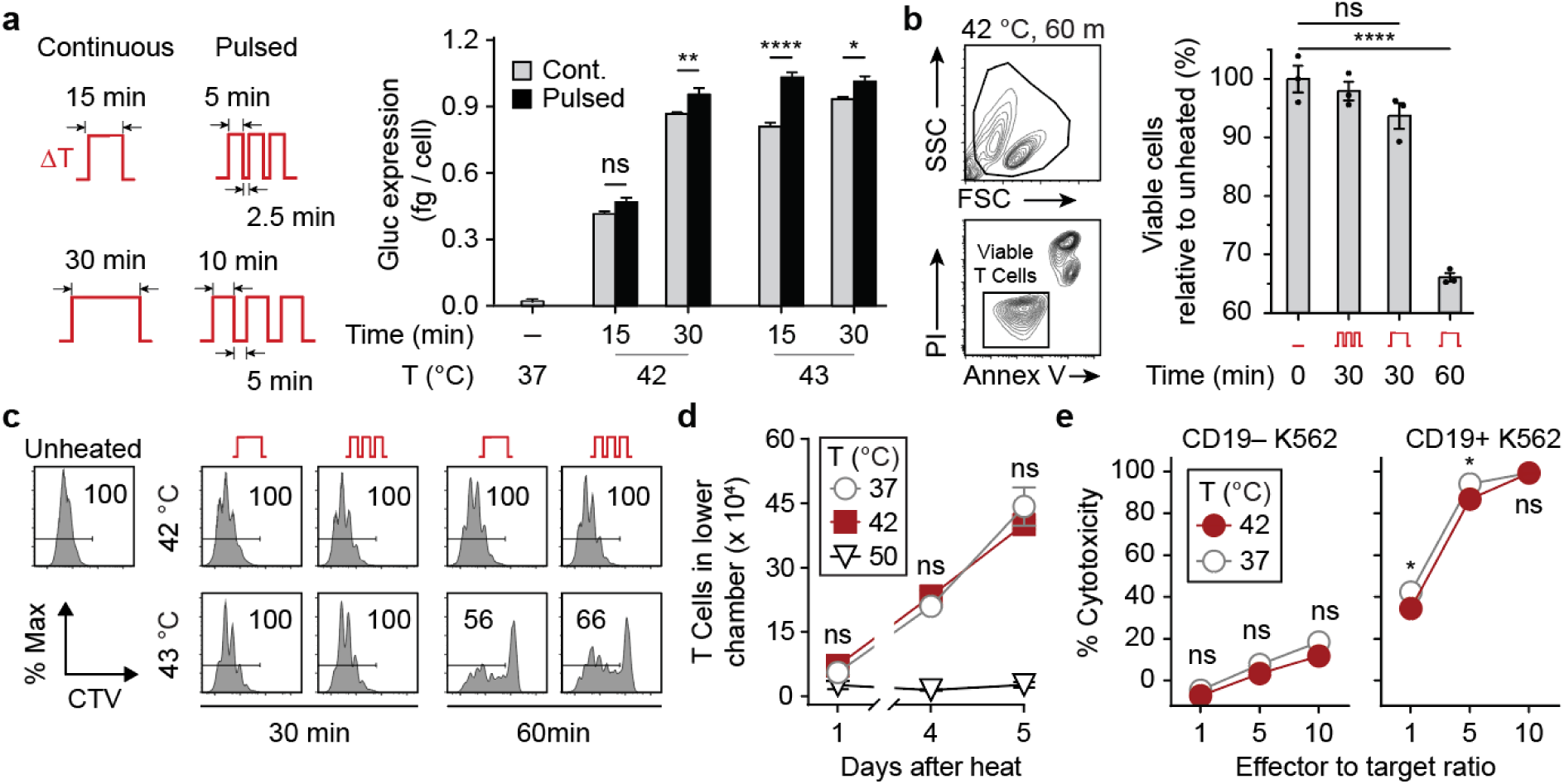
Thermal treatments are well-tolerated by primary human T cells. (**a**) Gluc activity of primary human T cells after continuous (grey) and pulsed (black) heat treatments with temperatures, total durations, and heating profiles as indicated (ns = not significant, *P<0.05, **P<0.01, ****P<0.0001, t-test, error bars show SEM, n = 3). (**b**) Propidium Iodide (PI) and Annexin V flow staining of CD3^+^ T cells. Bars represent viable populations (PI^−^Annexin V^−^) normalized to unheated samples (ns = not significant, ****P<0.0001, one-way ANOVA and Dunnett post-test and correction, error bars show SEM, n = 3). (**c**) CellTrace Violet (CTV) flow histograms of T cells after heat treatments and incubation with CD3/28 beads at a 3:1 bead to T cell ratio. (**d**) Number of cells in lower well of a transwell plate containing CXCL12. T cells were heated and loaded into the top well prior to sampling at indicated timepoints (ns = not significant between 37 °C and 42 °C, two-way ANOVA and Tukey post-test and correction, error bars show SEM, n = 3). (**e**) Percent cytotoxicity observed in CD19– or CD19+ luciferized K562 cells after incubation with T cells constitutively expressing CARs after heating with effector to target ratios as indicated (ns = not significant, *P<0.05, two-way ANOVA and Sidak post-test and correction, error bars show SEM, n = 3).

To probe the effects of thermal treatments on T cell migration, we used transwell assays to assess T cell chemotaxis and observed that heat treatments (42 °C for 30 min) did not significantly affect T cell migration into lower wells containing the chemokine CXCL12 compared to unheated controls whereas T cells heated to 50 °C were affected (**Fig. 2d**). To test longitudinal activation, we re-heated T cells over the course of 8 days and observed similar increases in GFP mean fluorescent intensity (MFI), as well as GFP activation and decay half-lives (t_1/2_ ∼0.5 and 1 day, respectively), suggesting that the magnitude and kinetics of T cell responses are unaffected by multiple heat treatments (**Fig. S3**). To quantify the effect of heat on T cell cytotoxicity, we incubated primary human T cells expressing an αCD19 CAR under a constitutive EF1α promoter with either CD19+ or CD19-K562s containing a luciferase reporter to allow quantification of cell death by loss of luminescence (**Fig. 2e**). At all effector to target cell ratios tested (1:1, 5:1, 10:1), heated T cells maintained greater than 90% of the cytotoxicity observed in unheated samples while no significant difference in cytotoxicity was observed in samples containing CD19-K562 target cells (**Fig. 2e**). Collectively, these data demonstrate that primary human T cells maintain the ability to proliferate, migrate, and kill target cells following short heat treatments delivered in continuous or pulsed wave forms for less than 30 minutes in duration.

### Thermal control of CAR T cell functions

The factors that contribute to durable anti-tumor responses include the degree to which key steps in the immunity cycle can be effectively directed against tumors^57^. Here we sought to build thermal circuitry to allow control over key CAR T functions including proliferation, antigen recognition, and T cell killing. We first explored thermal control of CAR T cell killing where the expression of an αCD19 CAR was placed under control of a thermal switch (TS-CAR) (**Fig. 3a**). Following heat treatment, TS-CAR T cells expressed αCD19 CARs to levels comparable to control T cells transduced with a constitutive vector (EF1α-CAR) (**Fig. 3b**). To test heat-triggered control of cytotoxicity, we heated TS-CAR T cells across a range of activation temperatures (40, 41, and 42 °C) and incubated them with either CD19– or CD19+ K562 target cells. We observed that intracellular Granzyme B levels – a cytotoxic effector molecule released by T cells – increased in CD19+ target cells as a function of temperature in contrast to control cells that lacked the CAR antigen (**Fig. 3c**). This was further supported by a TS-CAR T cell killing assay where we observed significant increases in CD19+ K562 cell death as the effector to target cell ratio was increased following thermal treatments at 42 °C (**Fig. 3d**).

**Figure 3.**
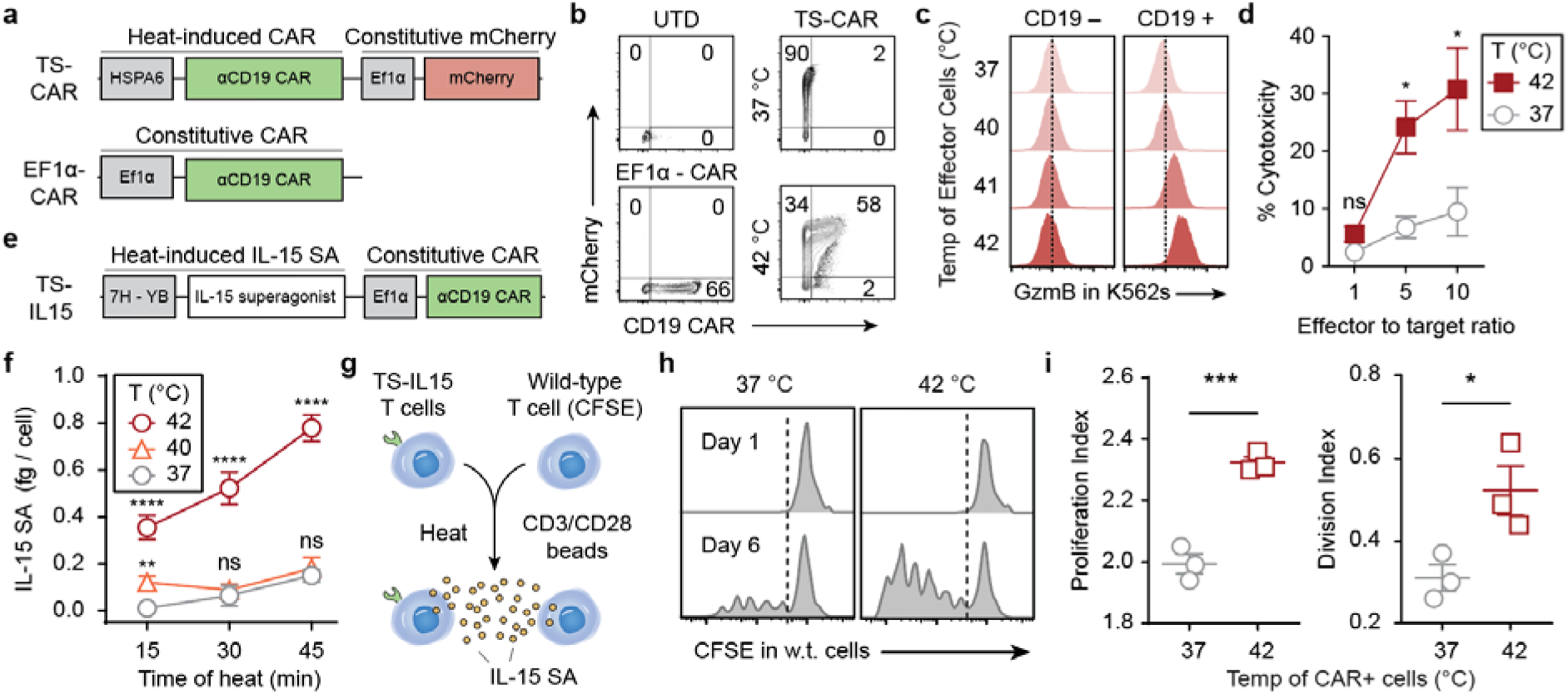
Thermal control of CAR T cell effector functions. (**a**) Schematic of a heat-inducible TS-CAR and constitutive EF1α CAR construct. Constitutive mCherry included in TS-CAR construct as a reporter for transduction (**b**) CAR flow staining with biotinylated CD19 on UTD, constitutive, and TS-CAR T cells. (**c**) Intracellular Granzyme B staining in K562 target cells after incubation with heated TS-CAR T cells. K562s lacking CAR antigen graphed in left column while CD19+ K562s are displayed in right column. (**d**) Killing of target CD19+ K562 cells by TS-CAR T cells (ns = not significant, *P<0.05, two-way ANOVA and Tukey post-test and correction, error bars show SEM, n = 3). (**e**) Schematic of TS-IL15 construct containing heat-triggered IL-15 superagonist and a constitutive CAR. (**f**) IL-15 superagonist concentrations in supernatant of TS-IL15 T cells following heat treatments. Temperature and duration of treatments are as indicated (ns = not significant, **P<0.01, ****P<0.0001, two-way ANOVA and Tukey post-test and correction, error bars show SEM, n = 3, comparisons are to unheated control). (**g**) Experimental design for co-culture assay of TS-IL15 cells and CFSE-labeled wild-type cells. CD3/28 beads were added at 1:10 bead to T cell ratio. (**h**) Representative flow histograms and (**i**) quantified proliferation and division indices as calculated by FlowJo proliferation tool of the CFSE-labeled wild-type T cell population (*P<0.05, t-test).

We further sought to determine whether thermal control could enhance T cell activation and proliferation. To do this, we cloned a single-chain IL-15 superagonist (IL-15 SA) comprised of the cytokine tethered to the sushi domain of the IL-15Rα subunit^58^ under control of our thermal vector (TS-IL15). IL-15 SAs are potent stimulants of CD8 T cells and NK cells and a clinical candidate, ALT-803, is currently under investigation for a wide range of cancers^59, 60^. To verify heat-triggered secretion of IL-15 SA, we analyzed conditioned media by ELISA and found that IL-15 SA levels increased with the duration and temperature of thermal treatment (**Fig. 3f**). To test whether heat-induced IL-15 SA was functionally active, we developed a T cell proliferation assay using CFSE-labeled wild-type T cells incubated with CD3/28 beads at a 10:1 ratio without supplemental cytokines. We found that this condition was insufficient to induce T cell proliferation compared to conditions when cytokines such as IL-2 was present in media (**Fig. S4**). Therefore, to test thermal control of IL-15 SA, we added heated or unheated TS-IL15 T cells to samples containing CFSE-labeled wild-type T cells with CD3/28 beads at a 10:1 T cell to bead ratio (**Fig. 3g**). Compared to unheated controls, we found that CFSE-labeled T cells in heated samples expanded with significantly higher proliferation and division indices (**Fig. 3h, i**), demonstrating that TS-IL15 T cells are capable of producing physiologically active levels of IL-15 SA following a single thermal treatment.

Last, we explored thermal control to expand CAR T recognition of tumor-associated antigens to include NKG2D ligands (NKG2DLs), which are upregulated on a wide range of cancers as well as suppressor cells^61-64^. Targeting NKG2DLs is limited by their expression in healthy cells such as intestinal epithelial cells and bone marrow stromal cells^65^. To enable thermal control of CAR T cells to target NKG2DL+ cells, we cloned a previously described NKG2DL-BiTE containing CD3-recognition domains from the OKT3 antibody linked to the extracellular domain of the human NKG2D receptor^64^. Our vector (TS-BiTE) included an Igκ leader sequence for BiTE secretion, a HisTag reporter, and a constitutive αCD19 CAR (**Fig. 4a**). After heat treatment, we observed that TS-BiTE T cells were stained positively on the cell surface by anti-HisTag antibodies compared to control cells transduced with a Fluc reporter vector (TS-Fluc) (**Fig. 4b**). Based on this, we postulated that T cells would activate by local BiTE binding to CD3 expressed by the same T cell (i.e., autocrine activation) before additional BiTEs would engage bystander T cells (i.e., paracrine activation). To test this, we heated a mixture of TS-BiTE transduced Jurkat T cells (i.e., CAR+) with untransduced cells (i.e., CAR-) as bystanders prior to co-incubation with NKG2DL+ CD19-K562 target cells (**Fig 4c–e**). This experimental setup allowed us to isolate T cell activation based on BiTE engagement without confounding factors due to CD19 CAR binding. We found that expression of the early activation marker CD69 on TS-BiTE Jurkat T cells was significantly upregulated compared to bystander cells as heating durations were extended (red versus black) (**Fig. 4f, g**). By contrast, CD69 was minimally upregulated on bystander cells compared to untransduced (UTD) Jurkat T cells that were incubated with K562 cells and heated in separate wells as controls (black versus gray). These data provided support that TS-BiTE T cells are primarily activated to target NKG2D ligands in an autocrine path. Finally, to quantify cytotoxicity from heat-triggered expression of BiTEs, we co-incubated primary human TS-BiTE T cells with NKG2DL+ CD19-K562 cells. In contrast to untransduced or TS-Fluc controls, TS-BiTE T cells secreted increasing levels of T_h_1 cytokines IFN-γ and TNF-α as temperatures were raised from 37 to 42 °C (**Fig. 4h**). We also observed temperature-dependent increases in K562 cytotoxicity but not at 37 °C compared to UTD controls, demonstrating lack of BiTE-induced killing at basal temperatures (**Fig. 4i**). Taken together, our data showed that thermal control can be extended to broad classes of immunomodulatory molecules to direct CAR T cell functions including proliferation, targeting, and cytotoxicity.

**Figure 4.**
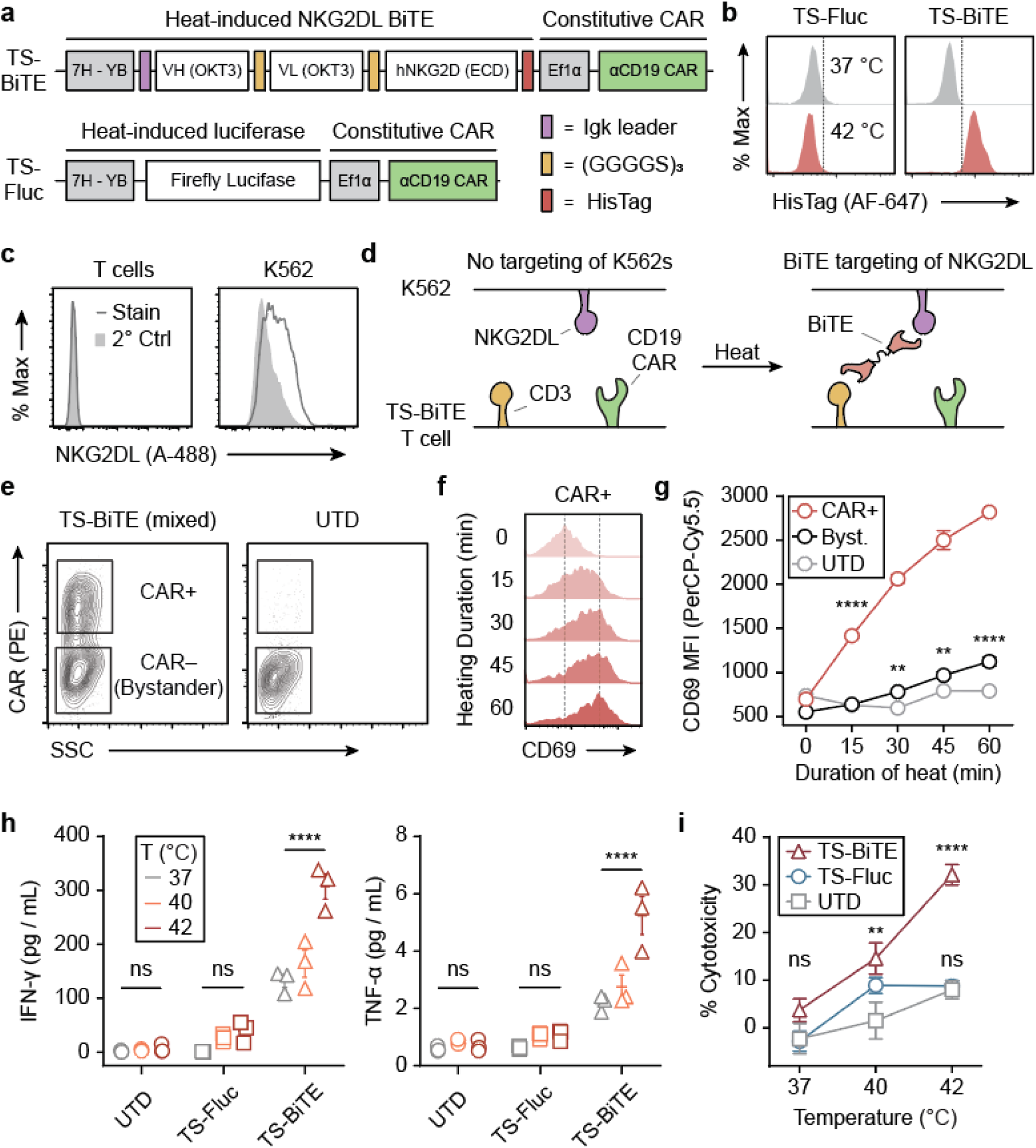
Expanding CAR T cell targeting via heat-triggered BiTEs. (**a**) Schematic of TS-BiTE and TS-Fluc thermal switches containing heat-triggered BiTE or Fluc reporters. Both constructs contained constitutive CARs. (**b**) Histograms for HisTag flow staining in TS-BiTE and TS-Fluc primary T cells following heating. (**c**) NKG2DL flow staining on primary human T cells and K562s using an NKG2D-Fc chimera and an αFc-A488 secondary antibody. Stain = full staining, 2° Ctrl = secondary antibody only. (**d**) Schematic depicting BiTE-mediated targeting of K562 target cells lacking the CAR target antigen via BiTE binding to NKG2DL and CD3. (**e**) Flow gating strategy for defining bystander cells based on CD19 CAR expression in Jurkat co-culture assays with K562s. UTD controls were gated on the lower (CAR-) population for graphing in (**g**). (**f**) Flow staining of CD69 on Jurkat T cells following heating and incubation with K562s. TS-BiTE CAR+ histograms (**f**) and summary data of indicated populations (**g**) are graphed (stats show comparison to UTD, ns = not significant, **P<0.01, ****P<0.0001, two-way ANOVA and Tukey post-test and correction, error bars show SEM, n = 3). (**h**) Cytokine concentrations in supernatant of primary human T cells after heat treatments and incubation with K562 cells. T cells were either untransduced or transduced with the indicated thermal switches (ns = not significant, ****P<0.0001, two-way ANOVA and Tukey post-test and correction, error bars show SEM, n = 3). (**i**) Cytotoxicity against K562s as quantified by luciferase assay after incubation with primary human T cells. T cells were either untransduced or transduced with the indicated thermal switches (ns = not significant, **P<0.01, ****P<0.0001, two-way ANOVA and Tukey post-test and correction, comparisons are to UTD control, error bars show SEM, n = 3).

### Photothermal targeting of CAR T cells enhances anti-tumor therapy

We next sought to determine whether thermal control of CAR T cells would enhance efficacy of adoptive therapies. To locally heat tumors, we used plasmonic gold nanorods (AuNRs) as antennas to convert incident near infra-red (NIR) light (∼650-900 nm) into heat^66^. PEG-coated AuNRs are well-studied nanomaterials with long circulation times that passively accumulate in tumors following intravenous administration^67, 68^. To confirm photothermal heating and thermal switch activation, primary T cells transduced with TS-Fluc were co-incubated with AuNRs in 96-well plates and irradiated with 808 nm laser light. In wells that reached 40–45 °C as monitored by a thermal camera, we observed a marked increase in luminescent signals when TS-Fluc T cells were present but not in wells containing untransduced controls (**Fig. 5a**), confirming plasmonic photothermal control of engineered T cells.

**Figure 5.**
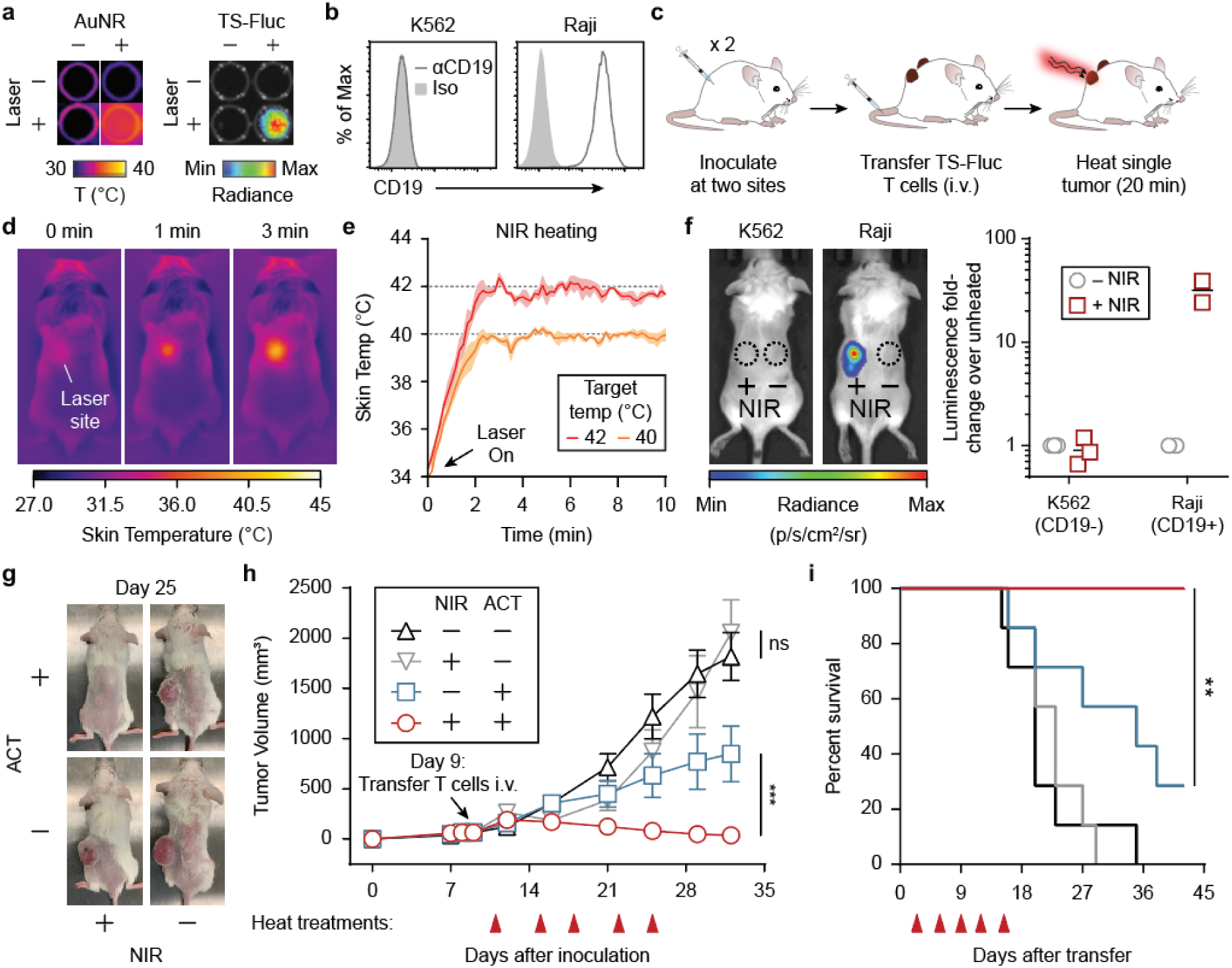
Photothermal targeting of CAR T cells improves anti-tumor responses. (**a**) Thermal and luminescent images of wells containing TS-Fluc T cells after irradiation with NIR laser light. Thermal images (left) were acquired using a FLIR thermal camera while luminescent images (right) were acquired using an IVIS Spectrum CT system (**b**) CD19 flow staining on K562 and Raji cell lines used for tumor inoculations, Iso = isotype control. (**c**) Schematic for experimental timeline where both flanks of each mouse were inoculated with the same cell line. TS-Fluc T cells were transferred via intravenous tail-vein injections and only one side of the mouse was heated. (**d**) Serial thermal images of tumor-bearing mouse during laser irradiation to heat tumor site. Timepoints are as indicated. (**e**) Thermal kinetic traces (colored lines) show average skin temperature of a 3 x 3 pixel ROI centered on laser site. Shaded regions around traces show standard deviation of 3 heating runs. (**f**) Left: Luminescent images of heated mice bearing either K562 (CD19-) or Raji (CD19+) tumors. Signal indicates luciferase activity by transferred TS-Fluc T cells. Right: Luminescence of each tumor site relative to the luminescence from the unheated tumor in the same animal. ROI’s were drawn as indicated in left panel. (**g**) Representative pictures of mice bearing CD19+ K562 tumors on Day 25. (**h**) Tumor growth curves following inoculation of CD19+ K562 T cells on Day 0, transfer of TS-IL15 T cells on Day 9 and heat treatments on Days 11, 15, 18, 22, and 25 (ns = not significant, ***P<0.001, two-way ANOVA and Tukey post-test and correction, error bars show SEM, n = 7). (**i**) Survival curves of tumor-bearing mice in (**g**) and (**h**) following transfer of TS-15 T cells and heat treatments (**p < .01, Log-rank (Mantel-Cox) test, n = 7).

To implement photothermal targeting in living mice, we inoculated NSG mice with bilateral flank tumors with one cohort receiving CD19– K562 cells and a separate cohort receiving CD19+ Raji cells to model CAR antigen positive and negative tumors (**Fig. 5b-c**). Following intravenous injection of AuNRs and adoptive transfer of T cells with constitutively expressed αCD19 CAR and thermal Fluc (TS-Fluc vector), we irradiated tumors with NIR laser light under the guidance of a thermal camera (**Fig. 5d**) to maintain target skin temperatures (**Fig. 5e**). After 20-minute heat treatments, luminescence was contained within Raji tumors receiving NIR light with Fluc activity upregulated greater than 30-fold at these sites compared to unheated Raji tumors in the same animal (**Fig. 5f**). Similar to *in vitro* experiments, CAR T cells could be repeatedly activated in Raji tumors (**Fig. S5**) but by contrast, we did not observe increased activity within antigen-negative K562 tumors following NIR heating (**Fig. 5f**). We attributed this lack of heat-induced activity to the greater than 20-fold lower density of intratumoral CAR T cells in resected K562 tumors that lack the CD19 CAR antigen (**Fig. S6**). Collectively these data demonstrate that the activity of intratumoral T cells engineered with thermal gene switches can be controlled by localized photothermal heating.

To augment CAR T cell therapies by thermal targeting, we adoptively transferred T cells transduced with TS-IL15 vector into NSG mice bearing CD19+ K562 tumors to allow thermal control of the single-chain IL-15 superagonist by cells constitutively expressing an αCD19 CAR. Photothermal heating of tumors was then carried out every 3-4 days (Days 11, 15, 18, 22, and 25) for a total of five treatments (**Fig. 5h**). Compared to control mice that did not receive CAR T cells or heat treatments (black), thermal treatment of tumor sites alone did not lead to reduction in tumor burden or improvement in survival (gray) (**Fig. 5g–i**). Transfer of TS-IL-15 CAR T cells alone significantly reduced tumor burden and improved survival (blue) yet greater than 70% (5/7) of animals reached euthanasia criteria within 38 days of ACT. By contrast, ACT of TS-IL15 CAR T cells combined with NIR treatments markedly reduced tumor burden and no animals reached euthanasia criteria within the time window of the study. Together, these data demonstrated that thermal targeting of tumors to control CAR T cell production of an IL-15 superagonist significantly improved tumor control and therapeutic outcomes.

## DISCUSSION

Here we developed a platform for remote thermal control of T cell activity. Remote control of T cells by small-molecules or light^69, 70^ either require systemic administration or are limited by light penetration through tissue. By contrast, thermal targeting of tissues can be accomplished by several platforms^25^ including focused ultrasound, which was recently demonstrated for control of T cell gene expression^71^ or thermal control of bacteria engineered with temperature-sensitive repressors^72^. To provide T cells the capacity to respond to heat, we designed synthetic thermal gene switches comprised of arrays of heat shock elements upstream of a core promoter. This architecture eliminated sensitivity to non-thermal stresses such as hypoxia and its thermal response was tunable based on the number of HSEs or different core promoters. While we tested constructs containing up to 7 HSEs paired with 4 core promoters, future work could explore a larger library of building parts including temperature-sensitive transcription factors such as HSF1 to tune response to heat. Importantly, we observed negligible activation of our thermal gene switches at temperatures ≤ 40 °C even when T cells were incubated for over 24 hours. This provides support that the temperature threshold for activation is higher than the range of typical fevers (∼38-40 °C)^73-75^ in patients with cytokine release syndrome (CRS), which would prevent T cell activation without a targeted thermal input. Despite this, we envision that the development of future thermal gene switches with lower temperature activation thresholds may be useful as a sense-and-respond circuit to autonomously detect fever temperatures and trigger expression of therapeutic agents such as tocilizumab to attenuate CRS.

With our current platform, we demonstrated thermal control of T cell activity using several classes of immunostimulatory genes including CARs, BiTEs, and cytokines. Engineered T cells that constitutively express similar classes of molecules have demonstrated strong anti-tumor efficacy but their therapeutic applications are limited by off-tumor effects and toxicities in healthy tissues^1, 2^. Thus, targeted expression of these genes within tumors could potentially contain potent T cell activity within the local TME and improve therapeutic outcomes. Given that the classes of molecules we studied included surface receptors, secreted cytokines, and bi-specific T cell engagers, we expect that a wide range of biologics are amenable for thermal control without potential loss of function due to protein misfolding or aggregation in T cells by heat stress. In the future, we anticipate that thermal targeting may have clinical use to treat cancer types that present as primary tumors with limited metastasis such as glioblastoma (GBM) or manage locally disseminated disease such as liver, lung, or brain oligometastases that are currently treated by surgical resection or ablation.

## Methods

### Plasmid construction

Synthetic thermal switches were produced as gene blocks by IDT and cloned into the Lego-C backbone (Addgene plasmid #27348). The core promoters were truncated immediately upstream of their previously described TATA boxes at their 5’-termini and at their translational start site on their 3’-termini ^76-78^. The genomic HSPA6 promoter was amplified from genomic DNA using PCR primers listed in a previous publication^56^. The NKG2DL BiTE sequence was described previously^64^ and modified to include an Igκ leader sequence to facilitate secretion from T cells as well as a HisTag for construct detection. This combined sequence was synthesized (ATUM) and cloned downstream of synthetic thermal switches. The IL-15 superagonist sequence was described previously^58^ and synthesized by ATUM without modification. The constitutive αCD19 CAR was kindly provided by Dr. Krishnendu Roy (Georgia Institute of Technology).

### Culture of primary human T cells and cell lines

CD19+ K562 (acquired from Dr. Yvonne Chen) and wild-type K562s (acquired from Dr. Krishnendu Roy) were cultured in Isocove’s Modified Dulbecco’s Medium (ThermoFisher #12440053) supplemented with 10% FBS (Fisher #16140071) and 10 U/ml Penicillin-Streptomycin (Life Technologies #15140-122). Raji cells were obtained from Dr. Krishnendu Roy and cultured in RPMI-1640 media supplemented with 10% FBS. Primary Human CD3+ cells were obtained from an anonymous donor blood after apheresis (AllCells) and were cryopreserved in 90% FBS and 10% DMSO until subsequent use. After thawing, cells were cultured in human T cell media comprised of X-VIVO 10 (Lonza #04-380Q), 5% human AB serum (Valley Biomedical #HP1022), 10 mM N-acetyl L-Cysteine (Sigma #A9165), and 55 µM 2-mercaptoethanol (Sigma #M3148-100ML) supplemented with 50 units/ mL human IL-2 (Sigma #11147528001).

### Lentiviral production and primary cell transduction

VSV-G pseudotyped lentivirus was produced via transfection of HEK 293T cells (ATCC) using psPAX2 (Addgene #12260) and pMD2.G (Addgene #12259); viral supernatant was concentrated using PEG-it virus precipitation solution (System Biosciences LV825A-1) according to manufacturer instructions. For viral transductions, primary human T cells were thawed, incubated for 24 hours, and activated with Human T-Activator Dynabeads (Life Technologies #11131D) at a 3:1 bead:cell ratio for 24 hours. To transduce the activated T cells, concentrated lentivirus was added to non-TC treated 6-well plates which were coated with retronectin (Takara #T100B) according to manufacturer’s instructions and spun at 1200 x g for 90 min at room temperature. Following centrifugation, viral solution was aspirated and 2 mL of human T cells (250,000 cells / mL) in human T cell media containing 100 units / mL hIL-2 were added and spun at 1200 x g for 60 min at 37 °C and moved to an incubator. Cells were incubated on a virus-coated plate for 24 hours prior to expansion and Dynabeads were removed 7 days after T cell activation. For cells flow-sorted prior to adoptive cell transfer, Dynabeads were added immediately after sorting at 3:1 ratios for 48 hours.

### Staining and flow cytometry

To detect CAR expression, biotinylated CD19 (10 µg / mL; Acro Biosystems #CD9-H8259) and Streptavidin-APC (ThermoFisher #S868) were used according to manufacturer instructions. NKG2DL expression was assessed by staining with NKG2D-Fc chimera (10 µg / mL; Fisher 1299NK050) followed by an αFc secondary stain (Invitrogen #A-10631). NIR Live/Dead (ThermoFisher #L34976), CFSE (LifeTech #C34554) and CellTrace Violet (CTV; LifeTech #C34557) were used according to manufacturer instructions. Human Fc block (BD #564220) was used prior to staining with any antibodies. For intracellular staining for Granzyme B, intracellular fixation and permeabilization buffers (eBioscience #88-8823-88) were used according to manufacturer instructions with Brefeldin A being added ∼4 hours prior to staining. Antibodies for Granzyme B (GB12; ThermoFisher), CD69 (FN50; BD), CD4 (RPA-T4; BioLegend); CD8 (RPA-T8; BioLegend), CD3 (UCHT1; BD), CD45 (HI30; BD), CD19 (HIB19; BioLegend), and HisTags (4E3D10H2/E3; ThermoFisher) were all used at 1:100 dilutions.

### *In vitro* luciferase and thermotolerance assays

Primary human T cells were heated in a thermal cycler and transferred to culture plates for incubation at 37°C. Unless otherwise noted, cellular supernatant was sampled for luciferase activity 24 hours after conclusion of thermal treatment. Non-thermal treatments were conducted by incubating engineered cells at indicated concentrations of CoCl_2_ (Sigma #232696-5G) or CdCl_2_ (Sigma #202908). When indicated, luminescence was compared to a ladder of recombinant Gaussia Luciferase (NanoLight #321-500) quantified using a Gaussia Luciferase Glow Assay Kit (ThermoFisher #16161) according to manufacturer’s instructions. For viability and proliferation studies, primary human T cells were heated in the thermal cycler prior to assaying with an apoptosis detection kit (BD # 556547) or CellTrace Violet (Fisher # C34571). For migration studies, wild-type cells were added to the top insert of a transwell plate (Sigma #CLS3421) while CXCL12 (50 ng / mL, Peprotech #300-28A) was added to the lower chamber. Cells in lower chamber were counted by hemocytometer at indicated times.

### Cytotoxicity and T cell activation assays

For cytometric analysis, TS-CAR T cells were heated in a thermal cycler and co-incubated with K562 target cells at a 10:1 effector cell to target cell ratio for 24 hrs prior to staining as described above. For luciferase-based assays, K562s were luciferized with either Firefly luciferase (CD19+) or Renilla luciferase (CD19-) and incubated with effector cells after heating. Unless otherwise noted, a 10:1 effector to target ratio was used. After incubation, either D-luciferin (Fisher #LUCK-2G; 150 µg / mL read concentration) or Rluc substrate (VWR # PAP1232; 17 µM read concentration). Maximum cytotoxicity was defined as luminescent signal from wells containing only media while no cytotoxicity was defined by wells containing only target cells. Supernatant was collected after incubation and assayed for cytokines using the human Th1/Th2/Th17 CBA kit (BD # 560484). IL-15 superagonist was quantified using the human IL-15/IL-15R alpha complex DuoSet ELISA (R&D Systems DY6924). For BiTE experiments with primary human T cells, two heat treatments (42 °C, 30 minutes) separated by 6 hours were applied to T cells prior to incubation with target cells.

### IL-15 superagonist Dynabead experiment

Wild-type primary human T cells were labeled with CFSE and incubated with either heated or unheated TS-IL15 cells. Beads were added at a 10:1 T cell to bead ratio that was determined not to induce strong proliferation in untransduced T cells without cytokine support (Fig. S4). CFSE labeling allowed discrimination from TS-IL15 cells and proliferation and division indices were calculated in FlowJo using the Proliferation tool.

### Animals

Eight-to sixteen-week old female NSG mice were used for all in vivo experiments. Mice were bred and housed in the Georgia Tech Physiological Research Laboratory (GT PRL) prior to start of experiments. All animal protocols were approved by Georgia Tech IACUC (protocols no. A100190 and 100191). All authors have complied with relevant ethical regulations while conducting this study.

### Photothermal heating and *in vivo* bioluminescence imaging

AuNRs were purchased from Nanopartz (# A12-10-808-CTAB-500) and pegylated (Laysam Bio # #MPEG-SH-5000-5g) to replace the CTAB coating. These AuNRs were intravenously injected into tumor-bearing mice (10 mg / kg) ∼48 hrs before adoptive transfer of T cells. Mice were anesthetized with isoflurane gas, and target sites were irradiated using an 808 nm laser (Coherent) under guidance of a thermal camera (FLIR model 450sc). Fluc activity was measured using an IVIS Spectrum CT (Perkin Elmer) ∼5 minutes after intravenous injections of D-luciferin (Fisher #LUCK-2G).

### Adoptive cell transfer (ACT) experiments

NSG mice were inoculated subcutaneously with 5 x 10^6^ Raji or K562 cell lines after the site was shaved and sterilized using an isopropyl wipe (GT PRL). ∼48 hrs prior to adoptive transfer of human T cells, pegylated AuNRs were injected intravenously via tail vein. Once tumors had reached ∼100-150 mm^3^, 1.5 × 10^6^ primary human T cells were injected via tail vein in 200 µL of sterile saline. Cells were transduced and sorted based on CD19 CAR expression prior to transfer as described above. After >24 hrs, photothermal heat treatments were administered and monitored as described above. Tumors were measured using digital calipers and volume calculated based on the equation length x width x depth x 0.52. Mice were euthanized when tumor volume exceeded 1500 mm^3^.

### Software and Statistical Analysis

All results are presented as mean, and error bars show SEM. Statistical analysis was performed using GraphPad Prism statistical software. For all graphs, * p < .05, ** p < .01, *** p<.001, **** p<.0001, ns = not significant. Flow cytometry data were analyzed using FlowJo X (FlowJo, LLC). Whole-mouse luminescence data were analyzed using Living Image software (PerkinElmer). Figures were designed in Adobe Illustrator.

## Data availability

All data supporting the findings of this study are available within the manuscript and its Supplementary Information. Raw data are available from the corresponding authors.

## Acknowledgements

We thank S.J. Priceman and J.M. Brockman for their helpful insights during planning and experiments. This work was funded by the NIH Director’s New Innovator Award (DP2HD091793), the National Center for Advancing Translational Sciences (UL1TR000454), the Shurl and Kay Curci Foundation. I.C.M. is supported by the Georgia Tech TI:GER program. L.G. is supported by the Alfred P. Sloan Foundation, the National Institutes of Health GT BioMAT Training Grant under Award Number 5T32EB006343 and the National Science Foundation Graduate Research Fellowship under Grant No. DGE-1451512. G.A.K. holds a Career Award at the Scientific Interface from the Burroughs Wellcome Fund. This work was performed in part at the Georgia Tech Institute for Electronics and Nanotechnology, a member of the National Nanotechnology Coordinated Infrastructure, which is supported by the National Science Foundation (Grant ECCS-1542174). This content is solely the responsibility of the authors and does not necessarily represent the official views of the National Institutes of Health.

## Author Contributions

I.C.M., and G.A.K. conceived of the idea. I.C.M., L.K.S., and G.A.K. designed the experiments. I.C.M. L.K.S., L.G., and A.Z. interpreted the results. I.C.M., L.K.S., and A.M.K. performed the experiments. I.C.M. and G.A.K. wrote the manuscript.

