## Supplemental Information for "Remote control of CAR T cell therapies by thermal targeting"

Address: Marcus Nanotechnology Building, 345 Ferst Drive, Atlanta, GA 30332, USA

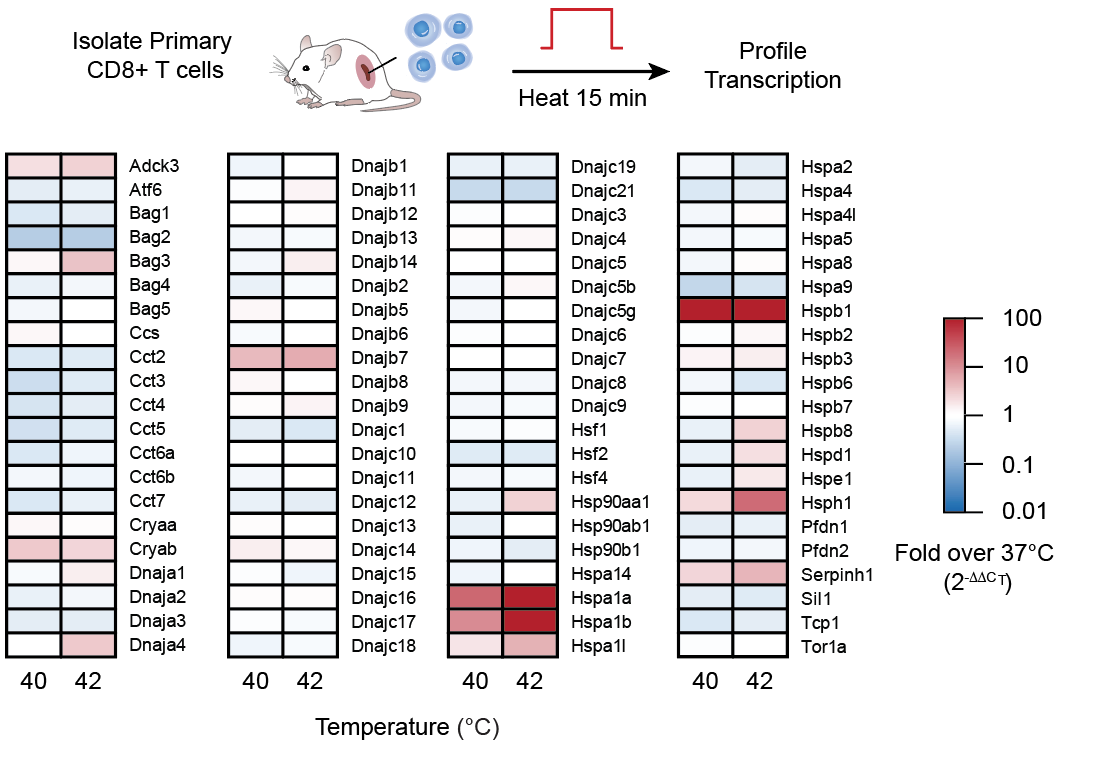

**Supplemental Figure 1. qPCR screen of HSPs in primary murine T cells.** Splenic CD8+ T cells were isolated using the CD8+ T cell isolation kit according to (Miltenyi 130-104-075). 6 hours after indicated heat treatments, mRNA was harvested and quantified using the Mouse HSP profiler kit (Qiagen PAMM-076Z) according to manufacturer instructions. Data are displayed relative to unheated controls.

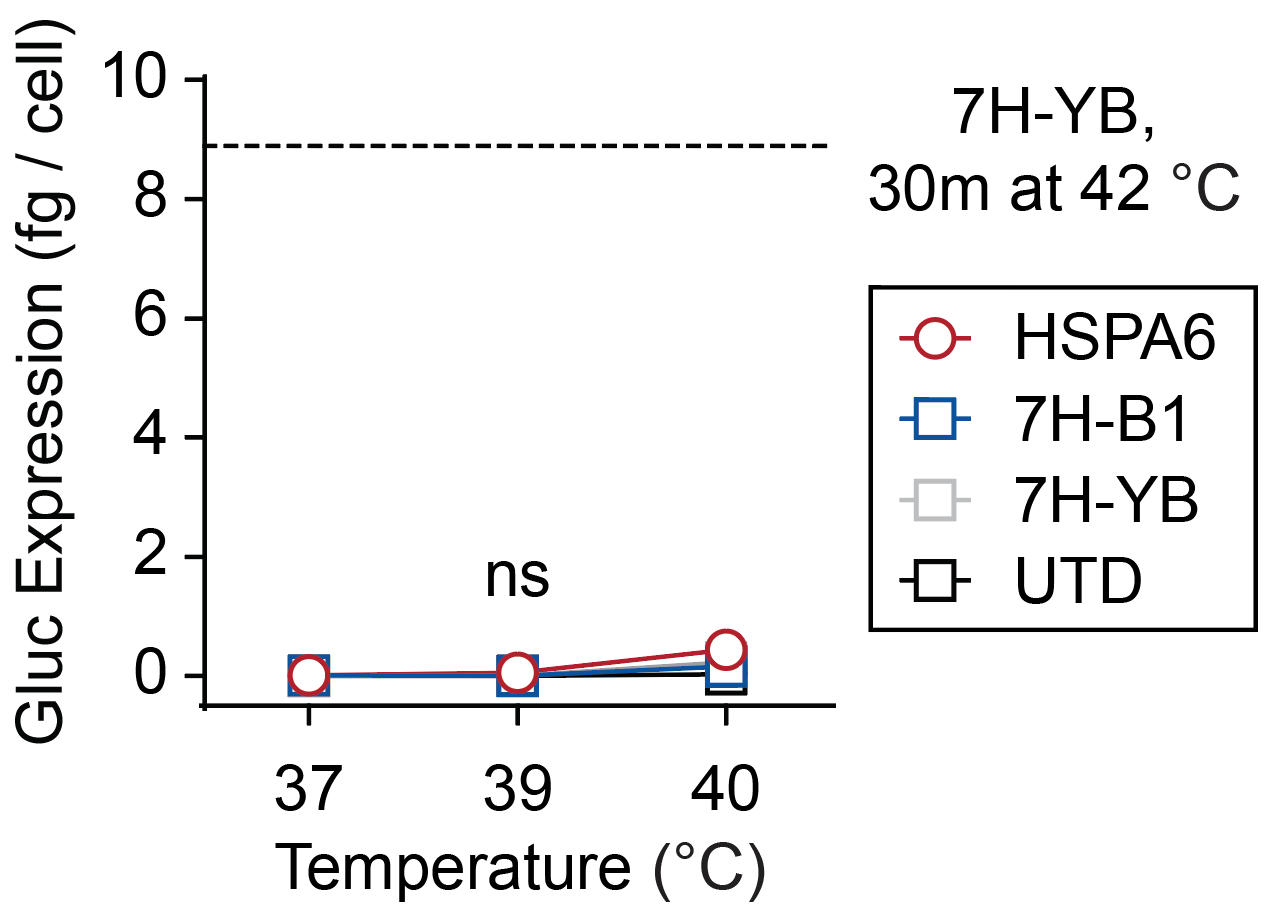

**Supplemental Figure 2: Prolonged heat treatments do not trigger thermal switches at lower temperatures.** Primary human T cells were transduced with the indicated thermal gene switches prior to incubation for 24 hours at the displayed temperatures, n = 3. For reference, dashed line indicates switch activity from the 7H-YB switch after a 42 °C treatment for 30 minutes (**Fig. 1i**).

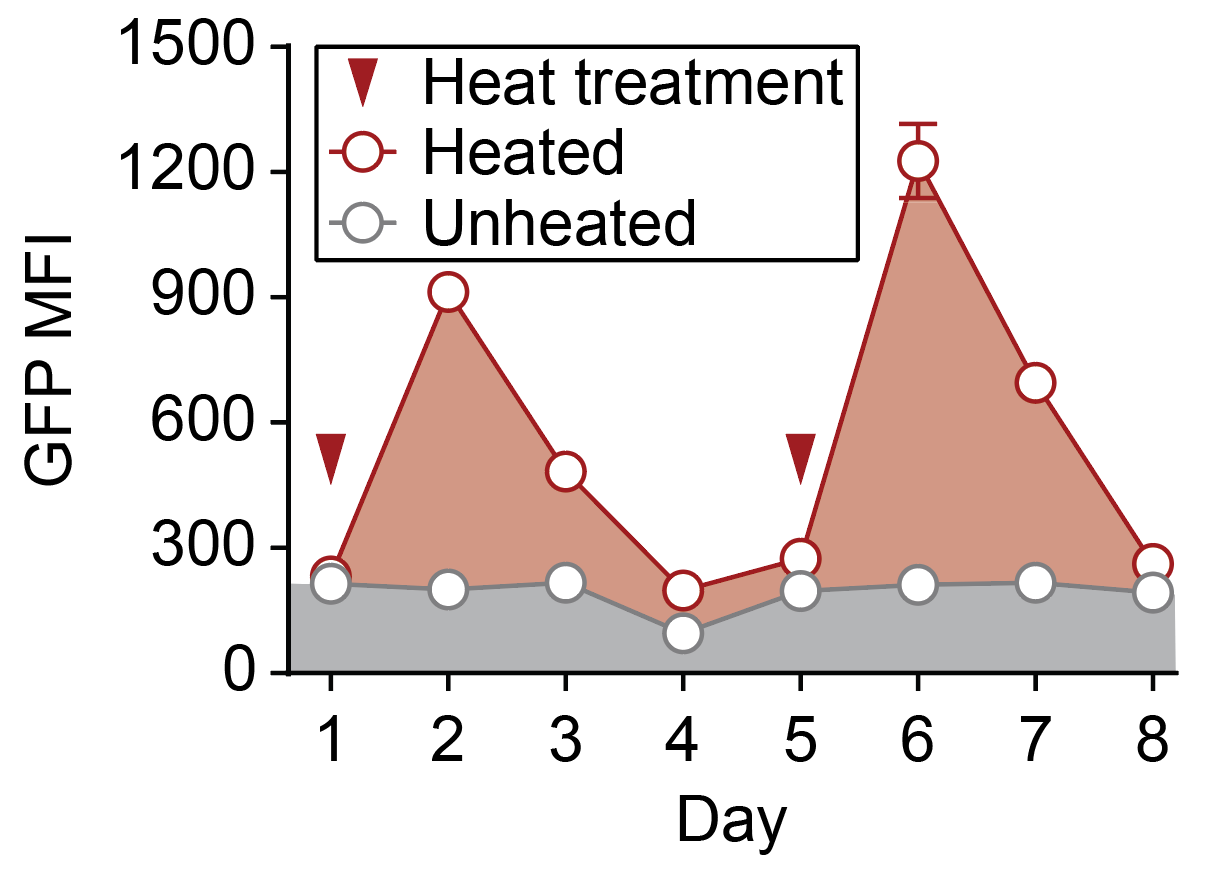

**Supplemental Figure 3: Longitudinal heating of primary human T cells.** Primary human T cells transduced with an HSPA6-GFP switch were repeatedly heated once GFP signal had returned to baseline after previous heat treatment. n = 3, error bars show SEM

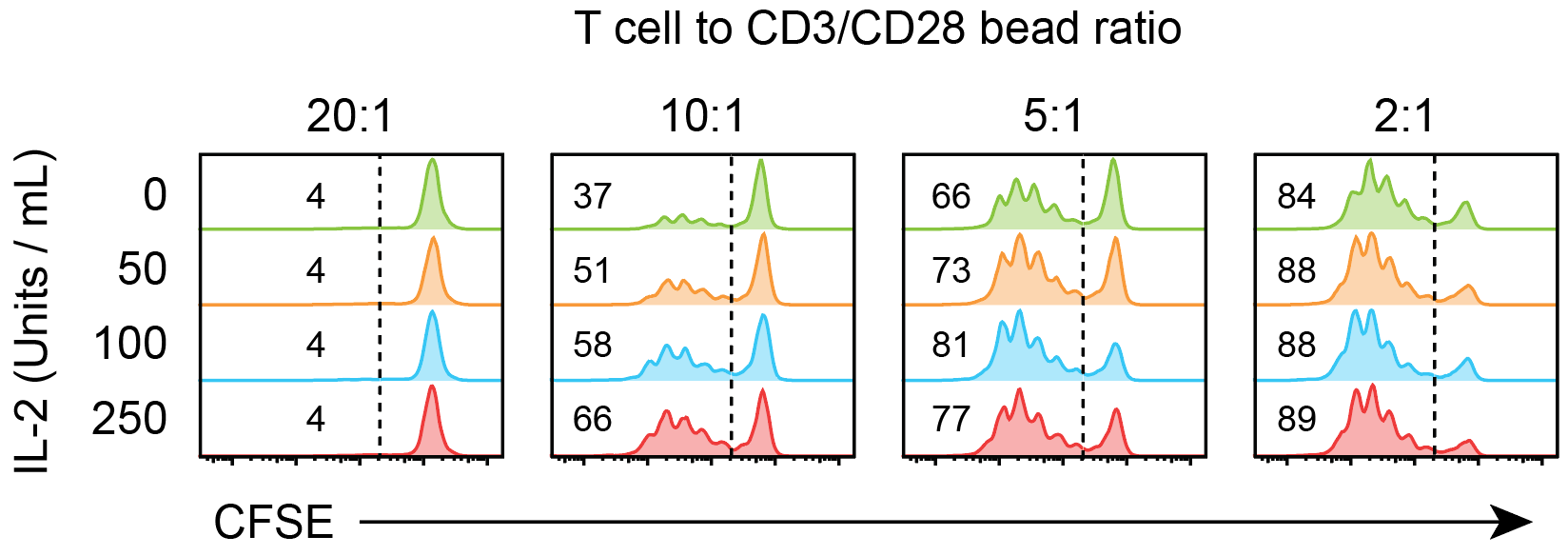

**Supplemental Figure 4: Cytokine support improves proliferation of T cells receiving low levels of CD3/28 stimulation.** T cells were labeled with CFSE and incubated with low levels of activating beads. For reference, routine expansion and culture of T cells uses 3 beads for every T cell. Increasing amounts of IL-2 were added to each bead ratio. All samples were assayed after 4 day incubations at indicated conditions.

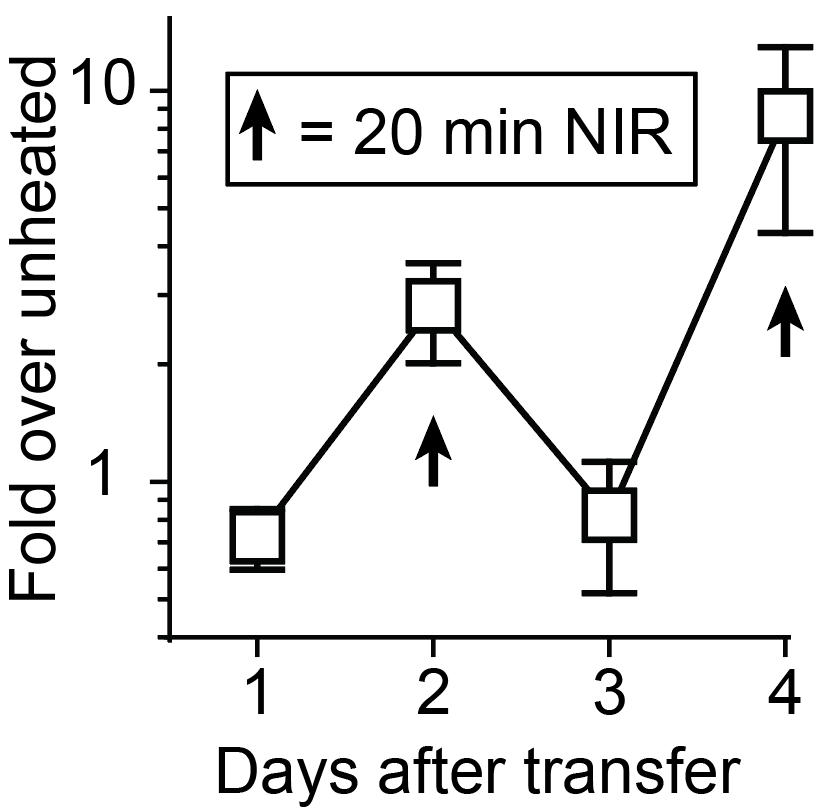

**Supplemental Figure 5: Longitudinal control of intratumoral CAR T cells using photothermal pulses.** Mice bearing Raji tumors (CD19+) were injected i.v. with TS-Fluc T cells. Tumor sites were irradiated on days 2 and 4 using NIR laser light as shown in **Figure 5d**. Luminescence was quantified daily via i.v. injections of D-luciferin, n = 3, error bars show SEM.

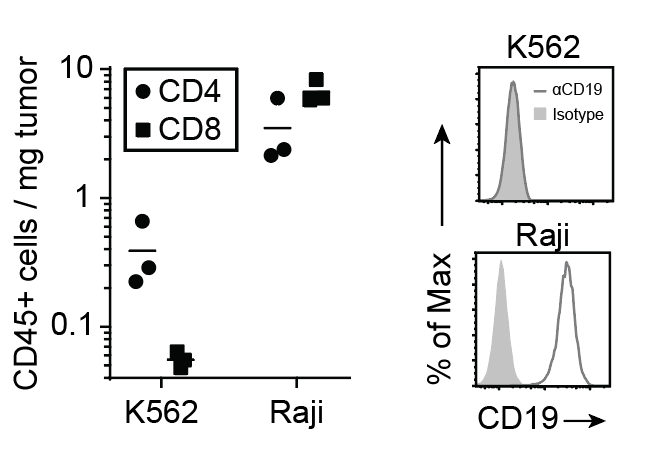

**Supplemental Figure 6: TS-Fluc CAR T cell infiltration into K562 and Raji flank tumors.** TS-Fluc T cells were injected i.v. into tumor bearing mice once tumors had reached ~250 mm^3^. After 7 days, tumors were resected, dissociated, and stained to quantify cellular infiltration using flow cytometry counting beads (ThermoFisher #C36950).
